# QTLViewer: An interactive webtool for genetic analysis in the Collaborative Cross and Diversity Outbred mouse populations

**DOI:** 10.1101/2022.03.15.484471

**Authors:** Matthew Vincent, Isabela Gerdes Gyuricza, Gregory R Keele, Daniel M Gatti, Mark P Keller, Karl W Broman, Gary A Churchill

**Author notes:** First-author (Matthew Vincent, Isabela Gerdes Gyuricza).

## Abstract

The Collaborative Cross (CC) and the Diversity Outbred (DO) mouse populations are related multiparental populations (MPPs), derived from the same eight isogenic founder strains. They carry >50M known genetic variants, which makes them ideal tools for mapping genetic loci that regulate phenotypes, including physiological and molecular traits. Mapping quantitative trait loci (QTLs) requires statistical and computational training, which can present a barrier to access for some researchers. The QTLViewer software is graphical user interface webtool for CC and DO studies that performs QTL mapping through the R/qtl2 package. Additionally, the QTLViewer website serves as a repository for published CC and DO studies, allowing users to explore QTL mapping results interactively and increasing the accessibility of these genetic resources to the broader scientific community.

## INTRODUCTION

Multiparental populations (MPPs) (de Koning and McIntyre 2017) improve upon traditional experimental crosses by interbreeding more than two isogenic strains capturing greater genetic diversity. The Collaborative Cross (CC) and Diversity Outbred (DO) mouse populations are related MPPs that have been used to genetically dissect complex traits, including obesity (Svenson et al. 2012), bone strength (Al-Barghouthi et al. 2021), short-term memory (Hsiao et al. 2020), and arsenic response (French et al. 2015). As recombinant populations, the CC and DO are well-suited for mapping quantitative trait loci (QTL), which can be powerful in studies of -omic traits, including gene expression, protein abundance, and chromatin accessibility (Abu Toamih Atamni et al. 2018; Aylor et al. 2011; Chick et al. 2016; Gerdes Gyuricza et al. 2022; Keele et al. 2020; Keele et al. 2021; Keller et al. 2018; Takemon et al. 2021).

The CC and the DO were bred from the same eight strains: A/J, C57BL/6J, 129S1/SvImJ, NOD/ShiLtJ, NZO/HlLtJ, CAST/EiJ, PWK/PhJ, and WSB/EiJ (Churchill et al. 2004; Churchill et al. 2012; Iraqi et al. 2012). The use of classical and wild-derived founder strains drives high levels of genetic variability (Yang et al. 2007; Yang et al. 2011). The CC represents ~60 recombinant strains that are fully inbred (>99%) whereas DO mice are outbred, thus each mouse is genetically unique. Mapping resolution is finer for the CC and the DO compared to intercrosses or backcrosses due to additional generations of meiosis, which facilitates the identification of candidate genes (Solberg Woods 2014).

Mouse experiments support the collection of multiple types of data on the same individuals, including physiological traits and molecular assays (*e.g*., gene expression) across multiple tissues. Integrative approaches like mediation analysis can be used to delineate the relationships among traits with shared QTLs (Chick et al. 2016; Keele et al. 2020). This series of analyses enable richer findings from CC and DO experiments but can also obstruct researchers without statistical and computational expertise.

Konganti et al. previously developed the gQTL webtool (Konganti et al. 2018), a graphical interface QTL mapping tool for CC data based on the R package DOQTL (Gatti et al. 2014). However, DOQTL is no longer maintained and instead the R/qtl2 package (Broman et al. 2019) should be used. Here, we present the QTLViewer software (https://qtlviewer.jax.org), an interactive QTL mapping webtool built for both CC and DO that utilizes the R/qtl2 package to perform QTL mapping as well as variant association mapping based on the founder strain genotypes. QTLViewer can perform mediation analysis for QTL through -omic traits to identify candidate causal mediators of physiological trait QTL. The QTLViewer website also serves as a repository for publicly available CC and DO datasets that can be downloaded for further investigation. This software will empower researchers to analyze and explore CC and DO data while facilitating access to relevant data and findings from others across the community.

## METHODS

The QTLViewer is an interactive web application designed to perform QTL mapping and related analyses in experimental CC or DO data. We make use of modern computing tools, such as application programming interfaces (APIs) and Docker containers, to make QTLViewer efficient, portable, and extendable.

### Implementation

The QTLViewer is comprised of three Docker (https://www.docker.com/ - v18.09.3) images that are managed with Docker-Compose (https://docs.docker.com/compose/ - v1.23.2). The user interface is contained in the churchilllab/qt2lweb container (v1.0.0), which is a Python (v3.6.9) web application that processes all requests made by the user. Each request is parsed and analyzed to determine which analyses need to be performed.

All computational requests are performed by the churchilllab/qtl2rest container (v0.1.0). This container loads data upon startup and listens to Web API calls via RestRserve (https://restrserve.org/ - v0.4.1). All data returned from these APIs are in JSON format. Since some requests require longer time to complete than others, we use another instance of the churchilllab/qtl2web container to manage them. We utilize Celery (https://docs.celeryproject.org/ - v4.47) with a Redis (https://redis.io/ - v6.2) backend as a task queue.

The containers work together to handle all requests. Requests generated from the web page are submitted via AJAX (Asynchronous JavaScript) and processed by the churchilllab/qtl2web container. A dialog box with a spinning wheel is displayed to the end user to show that the request is being processed. The request is parsed to determine which APIs need to be called. There could be one to many API calls per request. Those API calls are bundled together and submitted to the task queue for processing. Once the task queue receives the request, a unique ID is generated and sent back to the web page. The web page parses the unique ID and begins polling the task queue until the task is complete or an error occurs. While the web page is polling the task queue for a status, the API calls are submitted to the churchilllab/qtl2rest container for computational analyses. When all API calls are complete, the task queue bundles the data together and marks the job as complete. The background polling that has been happening on the web page, see that the job is complete, removes the dialog box, and processes the JSON data to display the appropriate graphs and data.

### Supporting Tools

QTLViewer relies on a set of tools that facilitate the analyses on the MPPs. Ensimpl (v 1.0.0) is a customized small version of Ensembl (https://www.ensembl.org) that provides a web API to retrieve genomic information. Ensimpl extracts the necessary data from Ensembl and creates a highly tuned SQLite (https://www.sqlite.org/) databases for querying purposes. The databases are separated by Ensembl release and species (mouse and human). The web API is written in Python utilizing FastAPI (https://fastapi.tiangolo.com/) framework. The API functionality provides search, genomic location lookup, gene lookup, and gene history.

In addition to Ensimpl, QTLViewer also utilizes a Founder SNP Database (https://churchilllab.jax.org/foundersnps). The Founder SNP database is a custom version of the Sanger SNP VCF files that have been processed via a Python script and stored in an SQLite (https://www.sqlite.org/) database. The SNP genomic location and allele calls for the eight founder strain alleles are stored. A custom Strain Distribution Pattern (SDP) is stored for easy query and dissemination of data. Currently GRCm38 (https://www.ncbi.nlm.nih.gov/assembly/GCF_000001635.20/) release 1410 and 1505 are supported.

### Statistical methods

The QTLViewer carries out several types of analysis to map and characterize QTL. QTL mapping analysis is run using the R/qtl2 package (Broman et al. 2019 – v0.28). The QTLViewer maps QTL by testing for an additive locus effect (additive QTL) or a locus-by-factor interaction effect (interactive QTL). For the additive model, the following linear mixed model is used to test an additive locus effect at loci spanning the genome:

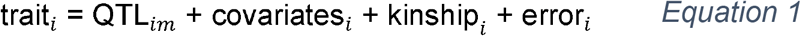

where trait_*i*_ is the phenotype value of mouse *i*, QTL_*im*_ is the effect of locus *m* on mouse *i* being tested, covariates_*i*_ is the cumulative effect of all covariates on mouse *i*, kinship_*i*_ is a random term that captures noise variation for mouse *i* due to population structure, and error_*i*_ is the independent random noise for mouse *i*. The structure of the kinship term is encoded in a genetic relationship matrix (K) estimated from the genotypes. We use the “leave one chromosome out” (LOCO) approach in which the K used for each locus *m* fit by Equation 1 excludes all markers from the chromosome of locus *m*, which improves QTL mapping power (Wei and Xu 2016).

For additive QTL, the QTL_*im*_ term represents allele dosages founder haplotypes at the locus. The QTLViewer will also plot the regression coefficients from the QTL_*im*_ term, *i.e*., the founder allele effects, when doing haplotype-based analysis. These effects can also be re-estimated as best linear unbiased predictors (BLUPs), which reduces the impact of rare alleles and can make signals clearer.

For interactive QTL, a similar model to Equation 1 is used to test a locus-by-factor interaction effect at loci across the genome:

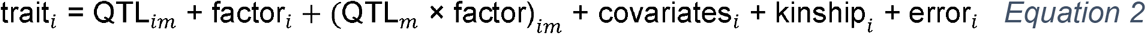

where factor_*i*_ is a covariate of interest for mouse *i* that may have an interaction effect with genotype at the locus *m*, (QTL_*m*_ × factor)_*im*_ is the QTL-by-factor interaction term at locus *m* being tested, and all other terms as defined before. Possible factors include sex and age.

The QTLViewer can also perform association mapping on bi-allelic variants using both additive and interactive models. Rather than encoding genetic effects based on doses of founder haplotypes, the QTL_*im*_ and (QTL_*m*_ × factor)_*im*_ terms in Equations 1 and 2 can be fit to doses of alleles of specific variants, imputed from dense genotypes of the founder strains. Although variant association mapping sacrifices some of the information present in the founder haplotypes, it does enable researchers to potentially identify specific variants of interests and prioritize candidate genes.

Finally, the QTLViewer can perform mediation analysis for additive QTL to identify candidate causal mediators (Chick et al. 2016; Keele et al. 2021; Keller et al. 2018) if -omic data, such as gene expression, have been collected on the same mice. Briefly, Equation 1 is re-used (with the kinship term excluded for computational efficiency) and the QTL of interest re-tested, but now conditioning one-by-one on candidate mediators. A strong candidate mediator will localize near the QTL and strongly reduce the QTL LOD score.

## RESULTS AND DISCUSSION

The QTLViewer webtool can be accessed at https://qtlviewer.jax.org/. After clicking “Datasets” on the top right of the screen (Figure 2A), users will find a list of publicly available datasets from CC and DO mice with their corresponding references and links to the specific QTLViewer instances (Figure 2B). Here we describe how a user would get started in analyzing a CC or DO dataset using a QTLViewer:

**Figure 1.**
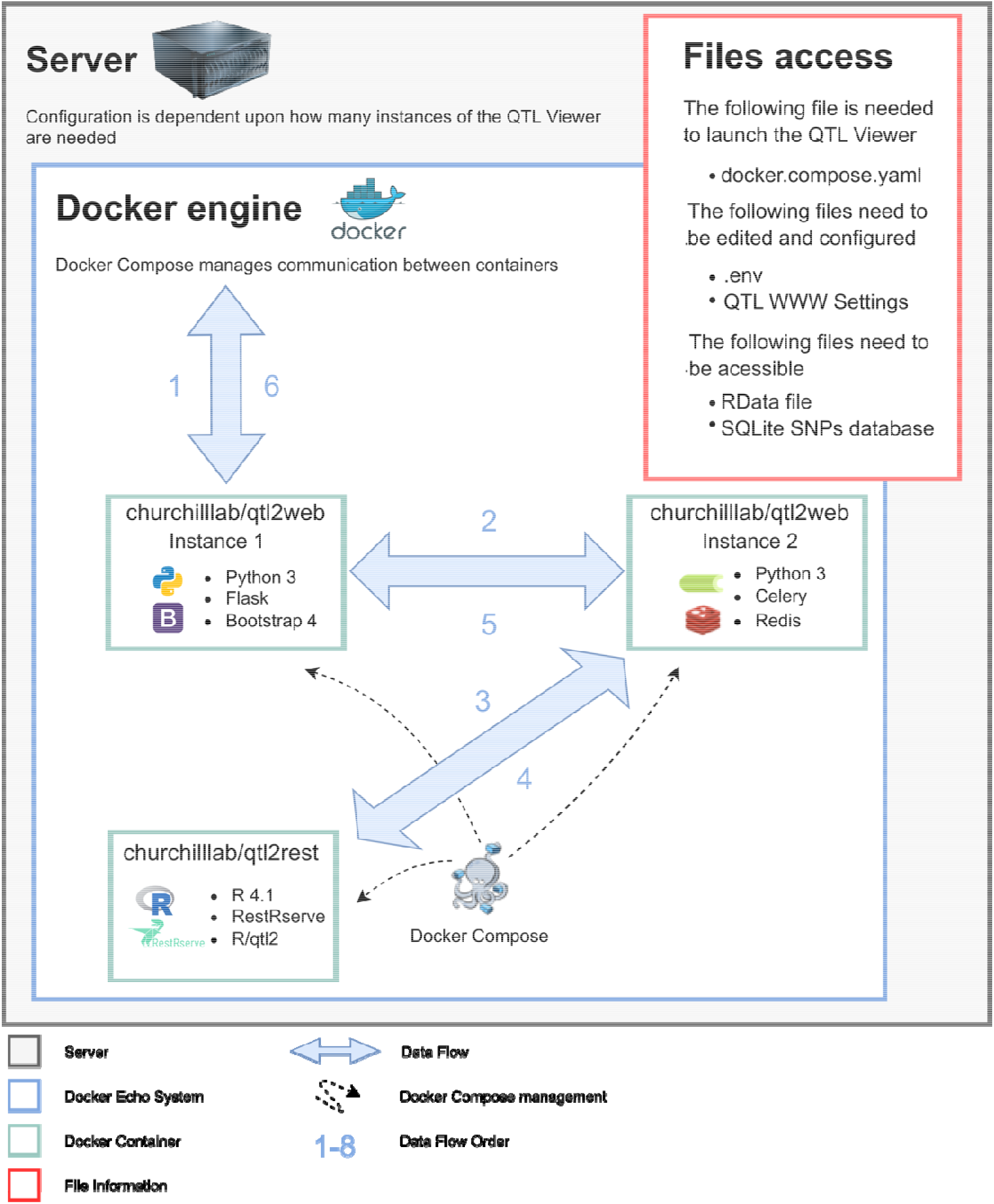
Diagram of QTL Viewer data flow. (1) URL is parsed and, if not an API call, returns requested information (Flask is used as the web application framework and Bootstrap 4 as the interface framework). (2) If the requested URL is an API call, the tool either returns a cached version if found or proceeds to call the API (Celery is used as a distributed task queue). (3) R/RestRserve package routes the URL to the correct R method to be performed. (4) A compressed JSON object is returned and if the HTTP request creates several QTL API calls, Redis will store the intermediate result from each call until all of them are finished. (5) Results are cached, and data is returned to Flask/Python. (6) The request is complete. The headers are checked and, if the response needs to be compressed, Flask will compress the data and send to the end user. The front-end user interface is now responsible for rendering the data.

**Figure 2.**
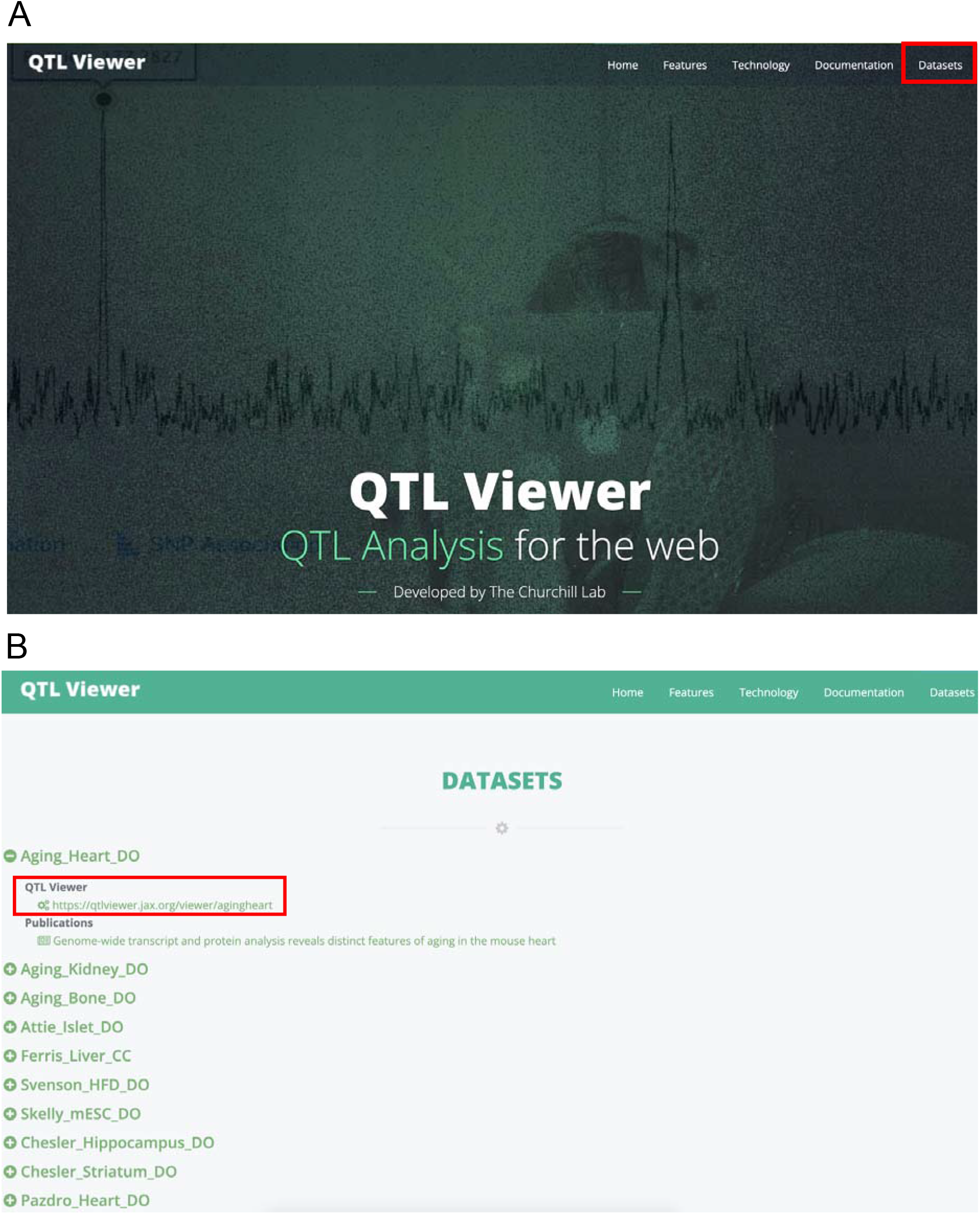
QTL Viewer access and public datasets. Users can access the main webpage of QTL Viewer at https://qtlviewer.jax.org/ where they will find a repository of different publicly available datasets (A). Clicking on “Datasets” reveals all available projects. Clicking on “+” button will show more information on the project of interest, including the QTL Viewer project link and related publications (B).

### Selecting a dataset

By clicking on the QTLViewer link corresponding to the project of interest, users will be directed to a webpage where they can explore the data. All QTLViewer projects contain at least one dataset such as a table of physiological traits, gene expression, or protein abundance data. Using the JAX Center for Aging Research DO data from the heart as an example (https://churchilllab.jax.org/qtlviewer/JAC/DOHeart) (Gerdes Gyuricza et al. 2022) we find transcript and protein data available. Users can switch datasets by clicking the selection box arrow next to the text “Current Data Set” (Figure 3A). All datasets in a QTLViewer instance are linked to a common set of mice and genotypes.

**Figure 3.**
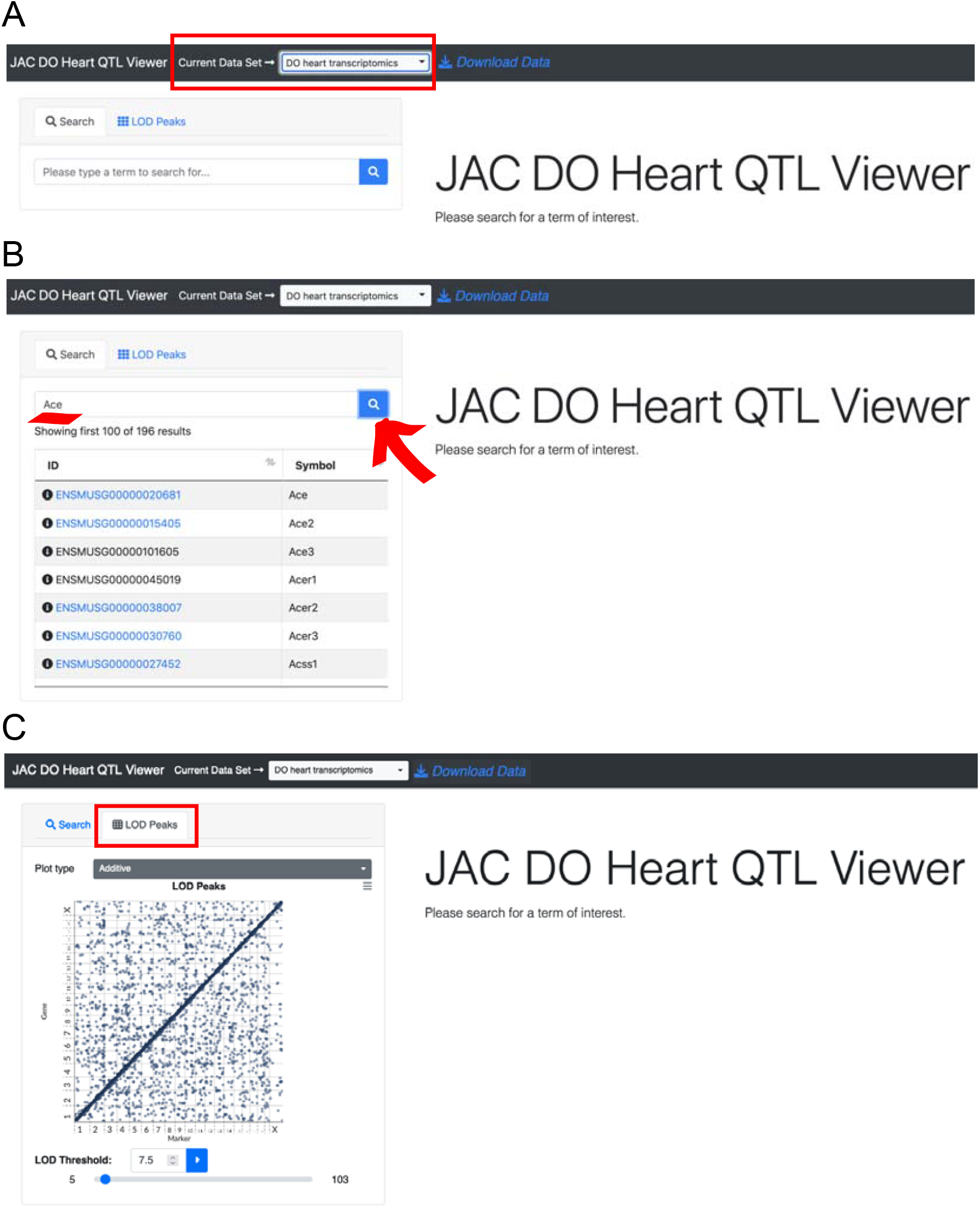
Navigating datasets in the QTL Viewer. The QTL Viewer page for a project can potentiall provides access to multiple datasets specific to the project which can be explored by changing the option on “Current Data Set”. In the screen shot, the aging DO heart transcriptomics data is selected (A). Within this specific dataset, it is possible to search for specific traits. For example, in the transcriptome dataset, the genes *Ace* and *Ace2* are available, but not *Ace3* (B). Switching to the “Lod Peaks” mode, visualizes a genome-wide transcriptome map with marker IDs on the x-axis and gene position on the y-axis, which is a common plot type to summarize all the expression QTLs (eQTLs) in the data according to a specified LOD threshold (C). This plot is particularly useful for visualizing genomic hotspots in the data, such as the one observed on chromosome 2.

### Searching the dataset by key word

QTL mapping and additional analyses can be performed for any trait in the data. To conveniently specify traits of interests, users can search via key word in the *Search* text box and press *Enter*. The search algorithm is specialized for -omics data like gene expression, recognizing gene symbols, gene names, and Ensembl identifiers. All elements that match the search criteria will be displayed in a table below, but only elements that are in the dataset will be displayed in blue and clickable (Figure 3B). This feature takes advantage of the Ensimpl database and webservice, which currently hosts human and mouse gene information with backward compatible versioning through Ensembl (Yates et al. 2020).

### Visualizing large-scale -omic QTLs (*e.g*., eQTLs)

The QTL mapping features of the QTLViewer are flexible and tailored to different types of data. For -omics data that have their own genomic coordinate, such as transcripts and proteins, a *LOD Peaks* tab is available next to the *Search* tab (Figure 3C). When selecting this option, a genomic grid is displayed with LOD scores from each gene or protein. This is the easiest way to look for regions of the genome where many QTLs co-map (genomic hotspots). The grid can be filtered based on a LOD threshold via a slider or manually setting the threshold in the *LOD Threshold* box. Factor-QTL interaction (for example, sex-interactive QTL) LOD peaks can be viewed by changing the *Plot type* select box. Additional information on specific QTLs can be interactively accessed by hovering and clicking on the points in the grid. The plot supports pan and zoom features.

### Generating genome-wide LOD plots (*i.e*., genome scans) for individual traits

Clicking on a point from the *LOD Peaks* grid or selecting a trait in the text search will generate a genome-wide LOD plot for the trait (Figure 4A). Information about each LOD score may be accessed by hovering. The *Plots* select box will switch between additive and factor-interactive QTL models. Clicking on a locus of interest (*i.e*., the peak LOD score) is used to drive additional analyses on the locus, which are visualized as plots in the lower panel. These analyses include estimation of founder allele effects, mediation of the locus effect through -omic traits, and SNP association mapping in the locus region.

**Figure 4.**
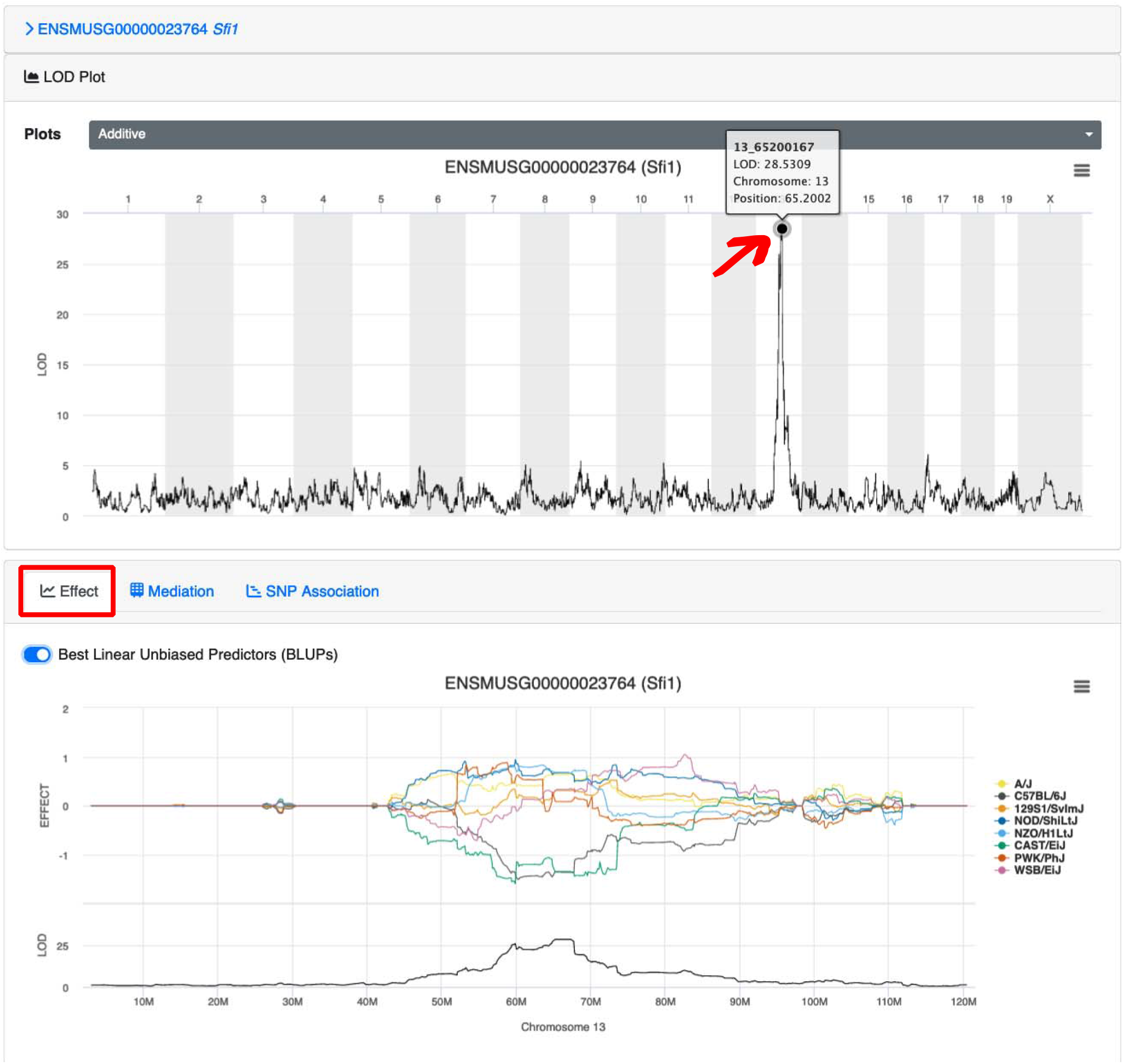
Genome-wide LOD and founder allele effects plots. Searching for specific traits or clicking on interesting QTLs on the transcriptome map reveals the genome-wide LOD plot with genome position across chromosomes on the x-axis and LOD scores on the y-axis. This plot reveals a strong eQTL on chromosome 13 at 65Mb for the gene *Sfi1*, with is a distant because *Sfi1* is encoded on chromosome 11. By clicking on the “Effect” function and then clicking back on the locus of interest produces a founder allele effects plot at the locus. The allele effects plot has genomic position in Mb on the x-axis and the estimated allele effects (top) and LOD scores (bottom) on the y-axis. The peak on chromosome 13 for the gene *Sfi1* is driven by lower expression from the CAST/EiJ and C57BL/6J alleles compared to higher expression from the other six alleles.

### Generating founder allele effects plot

The founder allele effects plot is the default analysis performed after clicking on a specific peak on the genome-wide LOD plot for a trait (Figure 4B). The allele effects plot shows the estimated founder allele effects across the chromosome that contains the selected peak. The allele effects can be reported as fixed effects coefficients (default) or as constrained Best Linear Unbiased Predictors (BLUPs) by checking the bar on top of the effect plot. BLUP estimates are generally preferable because they shrink extreme effects from rarely observed alleles, but they are computationally more intensive to calculate. These effects can be used to distinguish which founder strains likely possess the causal genetic variants. They also highlight QTL that are multi-allelic (Crouse et al. 2020) – a unique feature of MPPs like the CC and DO.

### Performing mediation of QTLs through -omic traits

Mediation analysis can be used to identify candidate mediators of a QTL for a trait. QTLViewer is primarily focused on using mediation analysis in the context of evaluating -omics traits as mediators. Mediation results and plots can be generated by clicking on the *Mediation* tab adjacent to the *Effect* tab, and then clicking back on the locus of interest on the LOD plot (Figure 5A). Doing so runs a mediation analysis which involves re-testing a QTL effect at the locus of interest, iteratively conditioning on candidate mediators. Promising candidates will significantly reduce or drop the initial QTL LOD score and be encoded on the genome near the QTL. This “LOD-drop” method of mediation is a powerful tool to identify candidate mediators, but it is also susceptible to false positive detection of linked independent effects (Chick et al. 2016; Crouse et al. 2020). Mediation can be performed against different datasets by changing the selection in the *Mediate Against* box.

**Figure 5.**
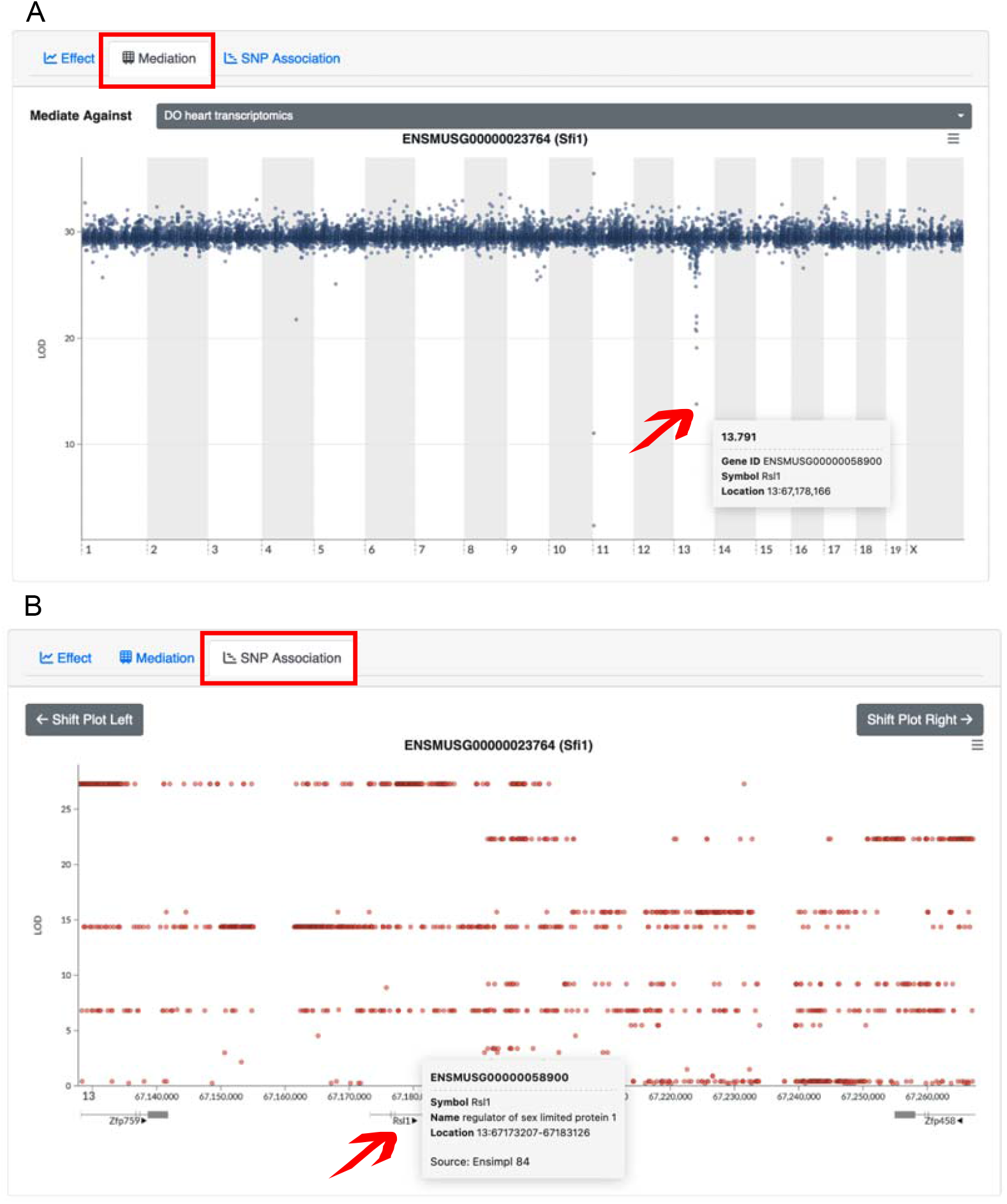
Mediation and SNP association plots. Mediation analysis can be performed on QTL of interest to identify candidate mediators as long as the QTL’s trait and the mediators are observed for the same mice. The mediation plot shows genomic position across chromosomes on the x-axis and conditional LOD scores on the y-axis. By mediating the distant eQTL of *Sfi1* on chromosome 13 through the DO heart transcriptomics data, the gene *Rsl1* on chromosome 13 at 67Mb is identified as a candidate mediator (A), which matches the eQTL position. A SNP association scan can be performed in the QTL region by clicking on the “SNP Association” button. The SNP association plot shows genomic position in bp on the x-axis and LOD scores on the y-axis. Annotated genes are overlayed below the x-axis. Variant association in the CC and DO reveals haplotypes shelves of variants in strong linkage disequilibrium (LD) with each other. Additional information on variants or genes can be accessed by hovering the cursor over the dot or gene track.

### Performing SNP association scans

The QTLViewer can perform SNP (or variant) association mapping in any selected region and can be useful to highlight genetic variants of interest in the QTL region. The SNP association plot can be generating by clicking the *SNP Association* tab adjacent to the *Mediation* tab, and then clicking on a locus of interest on the genome-wide LOD plot (Figure 5B). The SNP association plot shows LOD scores against genomic position for all variants overlaid with annotated genes by their corresponding genomic positions on the bottom. Users can pan and zoom in the SNP association plots. Hovering over a variant will display additional information such as variant consequences, and the allele distribution pattern among the founder strains.

### Exploring the relationships among traits and covariates

In addition to QTL mapping results, the QTLViewer can be used to explore the relationship between a trait and a covariate of interests (*e.g*., sex), which can be visualized on the bottom left corner of the screen, under the *Profile Plot* tab (Figure 6A). The plot can change based upon the covariates observed in the experiment by clicking on the *Select your factors* selection box, which will turn factors on or off. In addition, relationships among traits can be assessed based on correlation. The *Correlation* tab adjacent to the *Profile Plot* tab will show the correlation of the element of interest with any other element in the data (Figure 6B). The tool will display a scatter plot of the focus trait with another selected trait. To do that, users should choose a dataset in the *Select Correlation Dataset* box, and then select the element of interest from the table (Figure 6C). The values of the elements can be adjusted by the covariates specified on the *Covariate Adjustment* box, and points can be colored by specific covariates determined on *Select a series to color*.

**Figure 6.**
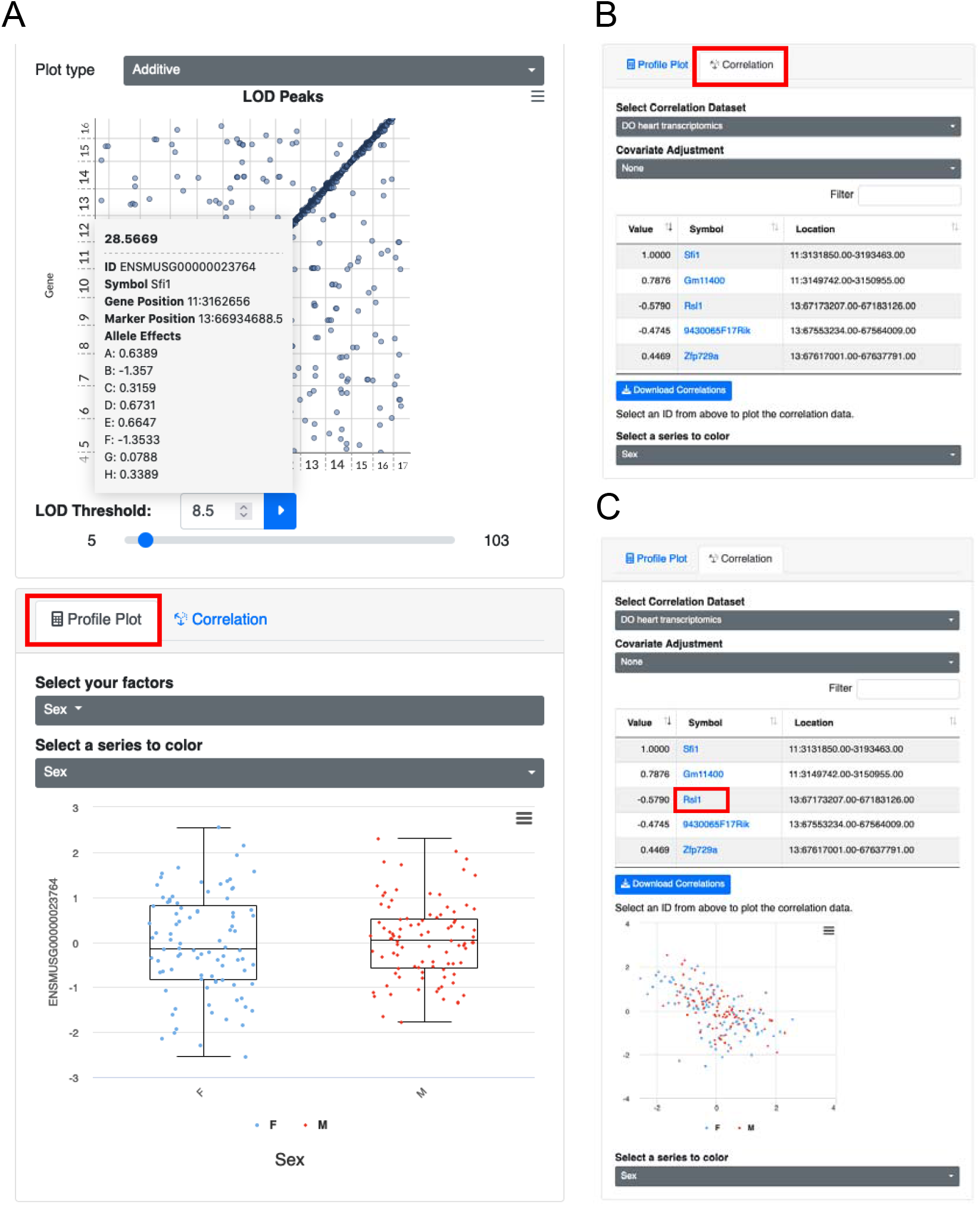
Plots for relating traits and covariates. After searching for a specific trait or selecting one based on its QTL, QTL Viewer can visualize the trait’s profile according to covariates in the data with “Profile Plot”. After clicking on the point corresponding to the gene *Sfi1* on the LOD peaks plots, QTL Viewer outputs the normalized expression of the *Sfi1* (y-axis) categorized by sex (x-axis) as boxplots (A). Additionally, QTL Viewer can display the correlation of a trait of interest with all the other elements of the data using the “Correlation” tab (B). The correlation data can also be downloaded locally by the user. All the elements on the correlation table are clickable. Clicking on “Rsl1” will generate a scatter plot between this gene and *Sfi1*, which illustrates the negative correlation between these two genes (C). *Sfi1* is negatively correlated with its mediator *Rsl1*, suggesting that *Rsl1* expression inhibits *Sfi1* expression.

### Downloading QTLViewer data objects

The QTLViewer contains multiple R data objects that supply all the necessary inputs for the analyses, including traits, covariates, and genotype probabilities. In addition to exploring the data interactively on the QTLViewer webpage, users can download the corresponding R data objects by clicking on *Download Data* on the top of the page (Figure 7A). All plots and analyses generated above can be downloaded as figures or data tables by clicking on the top right corner button on each plot (Figure 7A). When clicking on *Download Data*, users will be redirected to a new page displaying all the downloadable RData files (Figure 7B). There, users will find a “core” RData file containing all the input needed for mapping, such as genotype probabilities and marker information. Users can also download the “dataset” RDS files that contain the trait data, sample and assay metadata annotations, and a summary of the QTL mapping results (Figure 7C). This functionality enables users to quickly gain access to processed data files to run further analyses.

**Figure 7.**
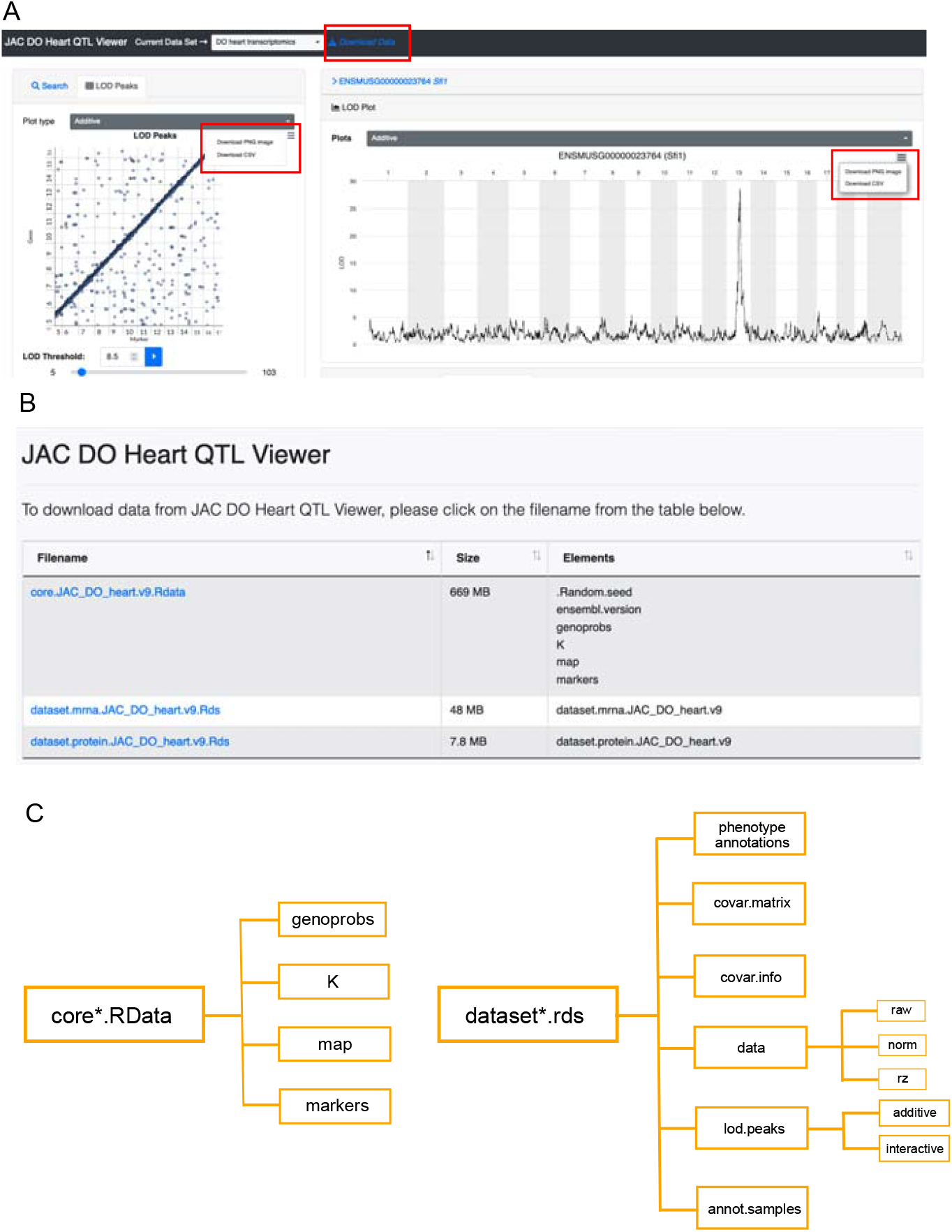
Figures and data download from QTL Viewer. All plots generated by QTL Viewer can be downloaded by clicking on the top right button in the application window (A). In addition to the figures, the processed data used as input to all analyses can be downloaded as R data files by clicking on “Download Data” (A). When using this option, users will be directed to a different webpage listing different components of the data for download (B). This includes a core RData object containing all information necessary for mapping, such as genotype probabilities, kinship matrix, and genomic map, and RDS files containing trait information, such as transcriptome (“dataset.mrna”) and proteome (“dataset.protein”) data. These RDS files are nested lists containing phenotype annotations, a matrix of covariates used for the QTL mapping (“covar.matrix”), information about the covariates (“covar.info”), trait data matrices, QTL mapping results (“lod.peaks”) and sample annotations (“annot.samples”). With gene expression data, “dataset.mrna” is a list with different forms of data as matrices, including the raw counts (raw) the normalized data (norm), and inverse normal transformed data (rz). The QTL mapping results “lod.peaks” is a list with QTL result tables from standard additive scans and potentially factor-interactive QTL scans.

### Future directions

The R/qtl2 software at the core of QTLViewer is very general and can accommodate a wide range of cross designs and data from model organisms other than the mouse. We plan to extend the QTLViewer to include mouse backcrosses, intercrosses, and two-way recombinant inbred panels. Adapting QTLViewer to MPPs from species other than the mouse will require modification of settings in the configuration files and the gene-search features for -omics data, and creation of a variant database.

We are in the process of adding a new method for mediation analysis based on Bayesian model selection (Crouse et al. 2021) that can provide richer inference for mediators of interest. We continue to expand the QTLViewer for new types of genomic data, such as chromatin profiling data, which will be incorporated and layered onto the genome browser tracks.

We continue to add new datasets to the QTLViewer website, which already represents a powerful resource for the community. We welcome contributions from outside investigators and will assist in the process of preparing their data for the QTLViewer.

## WEB RESOURCES

Users can access the QTLViewer webpage at https://qtlviewer.jax.org/. The QTLViewer software is version controlled and available from GitHub (https://github.com/churchill-lab). Installation instructions can be found in the Supplemental File.

The supporting tools Ensimpl and SNPDB are available online at https://churchilllab.jax.org/ensimpl and https://churchilllab.jax.org/foundersnps, respectively. The Docker containers code can be found at https://github.com/churchill-lab/qtl2web, https://github.com/churchill-lab/qtl2rest and https://github.com/churchill-lab/ensimpl. The Docker images are available at https://hub.docker.com/r/churchilllab/qtl2web/tags, https://hub.docker.com/r/churchilllab/qtl2rest/tags, and https://hub.docker.com/r/churchilllab/ensimpl/tags. The source code for QTL mapping analysis through R/qtl2 package can be found at https://github.com/rqtl/qtl2.

## DATA AVAILABILITY STATEMENT

The aging heart data analyzed here to demonstrate the QTLViewer functions can be found online at https://churchilllab.jax.org/qtlviewer/JAC/DOHeart and at Figshare under accession number 10.6084/m9.figshare.12378077.

