## Supplemental Table 1 for "QTLViewer: An interactive webtool for genetic analysis in the Collaborative Cross and Diversity Outbred mouse populations"

**QTLViewer setup**

*Prerequisite: Docker and Docker Compose installed*

To create your own QTL Viewer, a few configurations are needed. First, users should create a directory “project” to store files. The “project” directory will have the following directory structure:

Project

- docker-compose.yml
- .env
- project.viewer.settings
- snps.db3
- cache/
- data/

The following is a description of each content:

**docker-compose.yml**

This file should not be edited. It utilizes an environment file called .env that is located in the same directory. The .env file is what needs to be edited.

Example: <ftp://ftp.jax.org/churchill-lab/qtlviewer/example/docker-compose.yml>

**data/**

This directory needs to have Rds or Rdata files that contain the projects data. This needs to conform to theqtl2api RData file format (see <https://github.com/churchill-lab/qtl2api/blob/main/DataElement.md>).

**QTL Viewer Settings file**

Contains variables for all static text on the web page. It can be used to change the language of the QTL Viewer.

Example: <ftp://ftp.jax.org/churchill-lab/qtlviewer/example/project.viewer.settings>

**Founder SNPS DB**

SNP database for the SNP Association mapping.

Example: <ftp://ftp.jax.org/churchill-lab/qtlviewer/example/foundersnps.1505.snps.db3>

**.env**

This file is a list of all variables that need to be changed. The values you will need to change are highlighted in red.

Default Example:

\# Unique name of project

> COMPOSE_PROJECT_NAME=project1

\# Versions

> DOCKER_QTL2REST_VERSION=0.1.0

> DOCKER_QTL2WEB_VERSION=1.0.0

\# Docker Container

> CONTAINER_FILE_QTL2WEB_SETTINGS=/app/qtlweb/viewer.settings

> CONTAINER_DIR_QTL2WEB_CACHE=/app/cache

> CONTAINER_CACHE_NAME=projectcache

\# Name for Downloaded RData File

> QTLAPI_RDATA=None

> PYTHONUNBUFFERED=true

\# Ports

> HOST_PORT_WEB=8000

> HOST_PORT_API=8001

> HOST_PORT_REDIS=8002

\# Local File Paths

> HOST_FILE_RDATA=/project/data

> HOST_FILE_SNPDB=/project/data/foundersnps.1505.snps.db3

> HOST_FILE_QTL2WEB_SETTINGS=/project/project.viewer.settings

> HOST_DIR_QTL2WEB_CACHE=/project/cache

The following is the meaning of each value:

COMPOSE_PROJECT_NAME → a unique project name used for Docker internally

DOCKER_QTL2REST_VERSION → the Docker QTL2REST version to use

DOCKER_QTL2WEB_VERSION → the Docker QTL2WEB version to use

CONTAINER_FILE_QTL2WEB_SETTINGS → set to default

CONTAINER_DIR_QTL2WEB_CACHE → set to default

QTLAPI_RDATA → set to “None” (for Legacy application)

PYTHONUNBUFFERED → set to “true”

HOST_PORT_WEB → WWW Port for external use

HOST_PORT_API → API Port for external use

HOST_PORT_REDIS → Redis Port for external use

HOST_FILE_RDATA → full path to data directory

HOST_FILE_SNPDB → full path to Founder SNP DB

HOST_FILE_QTL2WEB_SETTINGS – full path to settings

HOST_DIR_QTL2WEB_CACHE → ull path to cache directory

CONTAINER_CACHE_NAME → unique name of cache

Example: <ftp://ftp.jax.org/churchill-lab/qtlviewer/example/.env>

Note: The only directory that needs “WRITE” access is /full/path/project1/cache.

Once users have edited the files, they should issue the following command on the command line to start the QTL Viewer:

> docker-compose up

This will download the necessary Docker images and start the server on the chosen port with the HOST_PORT_WEB value. Then, users should launch a browser to the name of the server (localhost for local installations) and the port: <http://localhost:8000>.
